# Flipping between Polycomb repressed and active transcriptional states introduces noise in gene expression

**DOI:** 10.1101/117267

**Authors:** Gozde Kar, Jong Kyoung Kim, Aleksandra A. Kolodziejczyk, Kedar Nath Natarajan, Elena Torlai Triglia, Borbala Mifsud, Sarah Elderkin, John C. Marioni, Ana Pombo, Sarah A. Teichmann

**Affiliations:** European Molecular Biology Laboratory-European Bioinformatics Institute (EMBL-EBI), Wellcome Trust Genome Campus, Hinxton, Cambridge, CB10 1SD, UK; Wellcome Trust Sanger Institute, Wellcome Trust Genome Campus, Hinxton, Cambridge, CB10 1SA, UK; Epigenetic Regulation and Chromatin Architecture Group, Berlin Institute for Medical Systems Biology, Max Delbrück Center for Molecular Medicine, Robert Roessle Strasse, 13125 Berlin-Buch, Germany.; Cancer Research UK London Research Institute, 44 Lincoln's Inn Fields, London, WC2A 3LY, UK; Department of Genetics, Evolution & Environment, University College London, Gower Street, London WC1E 6BT, UK; William Harvey Research Institute, Queen Mary University London, Charterhouse Square, London EC1M 6BQ, UK; Nuclear Dynamics Programme, The Babraham Institute, Babraham Research Campus, Cambridge, CB22 3 AT, UK; Cancer Research UK Cambridge Institute, University of Cambridge, Li Ka Shing Centre, Robinson Way, Cambridge CB2 ORE, UK

## Abstract

Polycomb repressive complexes (PRCs) are important histone modifiers, which silence gene expression, yet there exists a subset of PRC-bound genes actively transcribed by RNA polymerase II (RNAPII). It is likely that the role of PRC is to dampen expression of these PRC-active genes. However, it is unclear how this flipping between chromatin states alters the kinetics of transcriptional burst size and frequency relative to genes with exclusively activating marks. To investigate this, we integrate histone modifications and RNAPII states derived from bulk ChIP-seq data with single-cell RNA-sequencing data. We find that PRC-active genes have a greater cell-to-cell variation in expression than active genes with the same mean expression levels, and validate these results by knockout experiments. We also show that PRC-active genes are clustered on chromosomes in both two and three dimensions, and interactions with active enhancers promote a stabilization of gene expression noise. These findings provide new insights into how chromatin regulation modulates stochastic gene expression and transcriptional bursting, with implications for regulation of pluripotency and development.

## Introduction

Embryonic stem cells (ESCs) are capable of self-renewing and differentiating into all somatic cell types^1,2^, and their homeostasis is maintained by epigenetic regulators^3^. In this context, polycomb repressive complexes (PRCs) are important histone modifiers, which play a fundamental role in maintaining the pluripotent state of ESCs by silencing important developmental regulators^4^. There are two major polycomb repressive complexes: PRC1, which monoubiquitinylates histone 2A lysine 119 (H2Aub1) via the ubiquitin ligase Ring1A/B; and PRC2, which catalyzes dimethylation and trimethylation of H3K27 (H3K27me2/3) via the histone methyltransferase Ezh1/2.

Recently, we discovered that a group of important signaling genes co-exists in active and Polycomb repressed states in mESCs^5^. During the transcription cycle, recruitment of histone modifiers or RNA processing factors is achieved through changing patterns of post-translational modifications of the carboxy-terminal domain (CTD) of RNAPII^6^. Phosphorylation of S5 residues (S5p) correlates with initiation, capping, and H3K4 histone methyltransferase (HMT) recruitment. S2 phosphorylation (S2p) correlates with elongation, splicing, polyadenylation, and H3K36 HMT recruitment. Phosphorylation of RNAPII on S5 but not on S2 is associated with Polycomb repression and poised transcription factories, while active factories are associated with phosphorylation on both residues^5,7,8^. S7 phosphorylation (S7p) marks the transition between S5p and S2p^9^, but its mechanistic role is unclear presently.

Our genome-wide analyses of RNAPII and Polycomb occupancy in mouse ESCs (mESCs) identified two major groups of PRC-targets: (1) repressed genes associated with PRCs and unproductive RNAPII (phosphorylated at S5 but lacking S2 phosphorylation; PRC-repressed) and (2) expressed genes bound by PRCs and active RNAPII (both S5p and S2p; PRC-active)^5^. Both types of genes are marked by H3K4me3 and H3K27me3, a state termed bivalency^1,10^ H3K4me3 correlates tightly with RNAPII-S5p^5^, a mark that does not distinguish PRC-Active and Polycomb-represssed states.

The role of PRCs in modulating the expression of PRC-active genes was shown by PRC1 conditional knockout. Sequential ChIP and single-cell imaging showed mutual exclusion of S2p and PRCs at PRC-active genes^5^, although PRCs were found to co-associate with S5p. This indicates that PRC-active genes acquire separate active and PRC-repressed chromatin states. It remains unclear whether these two states occur in different cells within a cell population, or within different alleles in the same cell^5^. This pattern of two distinct chromatin states could imply a digital switch between actively transcribing and repressed promoters within a population of cells, thereby introducing more cell-to-cell variation in gene expression compared to genes with both alleles in active chromatin states.

Motivated by this hypothesis, here, we integrate states of histone and RNAPII modification from a published classification of ChIP-Seq data^5^ with single-cell RNA-sequencing data generated for this analysis. The matched chromatin and scRNA-seq data sets allow us to decipher, on a genome-wide scale, how differences in chromatin state can affect transcriptional kinetics. A schematic overview of our analysis strategy is shown in **Figure 1**. We focus on active PRC-target genes that are marked by PRCs (H3K27me3 modification or both H3K27me3 and H2Aub1) and active RNAPII (S5pS7pS2p), and compare these with “active” genes (marked by S5p, S7p, S2p without H3K27me3 and H2Aub1 marks). We quantify variation in gene expression and transcriptional kinetics statistically and by mathematical modeling (**Figure 1**). In addition, we map the functions of PRC-active genes in the context of pluripotency signaling and homeostasis networks. Further, we analyze the linear ordering and three dimensional contacts of PRC-active genes on the mouse chromosomes. Finally, we investigate the effect of Polycomb on regulating transcriptional heterogeneity by deletion of Ring1A/B followed by single-cell profiling.

**Figure 1.**
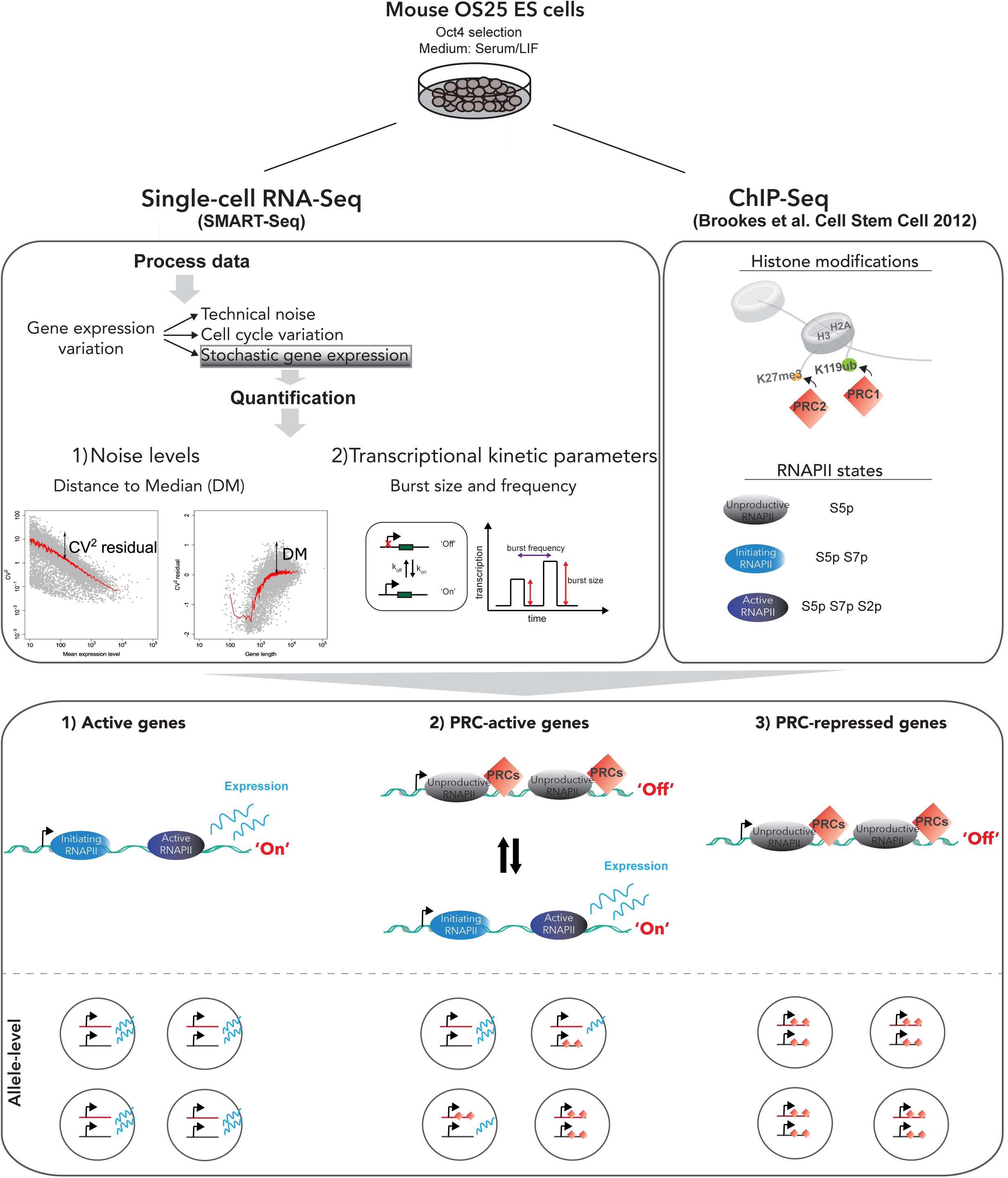
Summary of methodology. OS25 mESCs were cultured and characterized by single cell RNA-sequencing using the Fluidigm C1 system, applying the SMARTer kit to obtain cDNA and the Nextera XT kit for Illumina library preparation. OS25 cells are grown in conditions that select for undifferentiated cells (high Oct4-expressing). Libraries from 96 cells were pooled and sequenced on four lanes of a HiSeq. After quality control analysis of cells, 90 cells out of 96 remained for further analysis. We first unraveled contributions of components of gene expression variation using the scLVM method^13^. Removing cell cycle variation and technical noise allowed us to focus on stochastic gene expression. Gene expression variation can be quantified by squared coefficient of variation (CV^2^) or “distance to median” (DM), which is a measure of noise independent of gene expression levels and gene length. To explore the transcriptional kinetics of OS25 ES cells, poisson-beta model^16^ was fitted to single-cell gene expression data leading to estimates of burst frequency and size. Next, histone and RNAPII promoter modifications were obtained from Brookes et al.^5^ and integrated with single-cell RNA-seq to investigate relationship between stochastic gene expression and epigenetics. Active genes with no PRC marks are usually in the ‘on’ state with high burst frequencies (k_on_), PRCr genes are mostly ‘off’ and PRC-active genes switch between ‘on’ and ‘off’ states very frequently. Considering the allele-level possibilities, at active genes, both alleles would be in an actively transcribing state. For PRCa genes, both alleles would be in an actively transcribing state, or both alleles would be in a silent PRC-marked state, or only one allele is in PRC-marked state, which, subsequently, would result in noisier gene expression. For PRC-repressed genes, both alleles are expected to be in a silent PRC-marked state.

## Results

### Single cell RNA-sequencing and data processing

To investigate how Polycomb repression relates to stochasticity in gene expression, we profiled single cell transcriptomes of mouse OS25 ESCs cultured in serum/LIF, previously used to map RNAPII phosphorylation and H2Aub1^5^. Single cell RNA-sequencing was performed using the Fluidigm C1 system, applying the SMARTer kit to obtain cDNA and the Nextera XT kit for Illumina library preparation. Libraries from 96 cells were pooled and sequenced on four lanes of an Illumina HiSeq2000 (**Figure 1**; please refer to Methods for details).

Next, we performed quality control analysis for each individual cell dataset and removed poor quality data based on two criteria (as described before in^11^). Cells were removed if: (1) the total number of reads mapping to exons for the cell was lower than half a million, (2) the percentage of reads mapping to mitochondrial-encoded RNAs was higher than 10%. We also compared normalized read counts of genes between cells and found many genes abnormally amplified for three cells. Therefore, we removed these cells, resulting in 90 cells that could be used for further analysis. For these 90 cells, over 80% of reads were mapped to the *Mus musculus* genome (GRCm38) and over 60% to exons (**Supplementary Fig. 1A-C**).

OS25 ES cells are grown under Oct4 selection and do not express early differentiation markers such as Gata4 and Gata6^5^, having the expected features of pluripotency. They are ideal for studying Polycomb repression and its impact on transcriptional cell-to-cell variation as compared to other culture conditions such as 2i (serum free). ESCs grown in 2i show decreased Polycomb repression and RNAPII poising at well characterized early developmental genes^12^, therefore making 2i conditions the least ideal conditions to study mechanisms of Polycomb regulation in the pluripotent state. As previously shown^5^, we do not observe distinct subpopulations of cells based on key pluripotency factors and differentiation markers in our OS25 single cell datasets (**Supplementary Fig. 1D**).

Additionally, we compared single cell expression profiles of the OS25 ESCs grown under Oct4 with recently published scRNAseq datasets from mESCs cultured in serum+LIF and 2i^11^, Principal component analysis using pluripotency genes and differentiation markers shows that OS25 cells are more similar to the subpopulation of pluripotent serum cells, rather than the subpopulation of serum cells that are either “primed for differentiation” or “on the differentiation path”. (**Supplementary Fig. 1E**).

### Defining chromatin state and gene expression noise for each gene

We integrated our new single-cell RNA-seq data with a previous classification of gene promoters according to the presence of histone and RNAPII modifications^5^ (**Figure 1**). Comparison of our average single-cell expression profiles with the bulk gene expression (mRNA-seq) profiles from Brookes et al.^5^ yields a high correlation (Spearman’s rho = 0.87, **Supplementary Fig. 1F**), suggesting that the chromatin and RNAPII data reflect cells in the same biological state as the single cell RNA-seq data.

Next, we analyzed gene expression variation within the single-cell data. First, we quantified cell-to-cell variation at each mean expression level using the coefficient of variation (**Supplementary Fig. 2A**). Cell-to-cell variation can arise either due to stochastic gene expression itself, or technical noise or confounding expression heterogeneity due to biological processes such as the cell cycle.

To isolate pure stochastic gene expression from cell cycle variation in gene expression, we applied a latent variable model^13^. This is a two-step approach, which reconstructs cell cycle state before using this information to obtain “corrected” gene expression levels. The method reveals that the cell cycle contribution to variation is 1.2% on average (**Supplementary Fig. 2B**). While this effect is small, when clustering all cells based on G2/M stage markers, we found that cells separate into two groups: one with high expression of G2 and M genes and the other with low expression of these genes (**Supplementary Fig. 2C**). Applying the cell cycle correction removes this effect, leading to a more homogeneous expression distribution of these genes across the cells (**Supplementary Fig. 2D**).

To account for the technical noise present in single cell RNA-seq data, we removed lowly expressed genes that are most likely to display high technical variability^14,15^. Here, a gene is considered as lowly expressed if the average normalized read count is less than 10. This results in a set of 11,861 genes with moderate to high mRNA abundance. Subsequently, we use the DM (distance to median) to quantify gene expression variation in mRNA expression^11^, since it accounts for confounding effects of expression level and gene length on variation (described in detail in the Methods; **Figure 1**).

Among the 11,861 expressed genes, 7,175 have categorized ChIP-seq profiles as defined by Brookes et al.^5^; genes excluded have TSS regions that overlap with other genes, and therefore cannot be unequivocally classified. We defined two major sets of genes based on their PRC marks and RNAPII states: (1) “Active” genes (n=4,483) without PRC marks (H3K27me3 or H2Aub1) but with active RNAPII (S5pS7pS2p), (2) “PRC-active” genes (labeled as “PRCa”; n=945) with PRC marks (H3K27me3 or H3K27me3 plus H2Aub1) and active RNAPII.

To explore the transcriptional kinetics of these genes and describe stochastic gene expression in OS25 ES cells, we estimated their kinetic transcription parameters using a Poisson-beta model described previously^16^ (see also in the Methods).

### PRCa genes have distinct transcriptional kinetics and noise profiles

Using the DM measure to quantify gene expression variation in single cells, we observe that histone modifications mediated by PRCs (H3K27me3 or H3K27me3&H2Aub1) correlate with high levels of variability compared to Active genes (those without PRC marks; *P* < 2.2×10^-16^ by the two-tailed Wilcoxon rank sum test, **Figure 2A**). Furthermore, the inferred kinetic parameters provide insight into the expression behavior of genes, showing that active genes have significantly higher burst frequencies than PRCa genes (**Figure 2A and Supplementary Fig. 3A**). This suggests that PRCa genes are more frequently in the “off” state, *i.e.* more alleles are in the off state at any given point in time, potentially due to the PRC repression of a subset of alleles.

**Figure 2.**
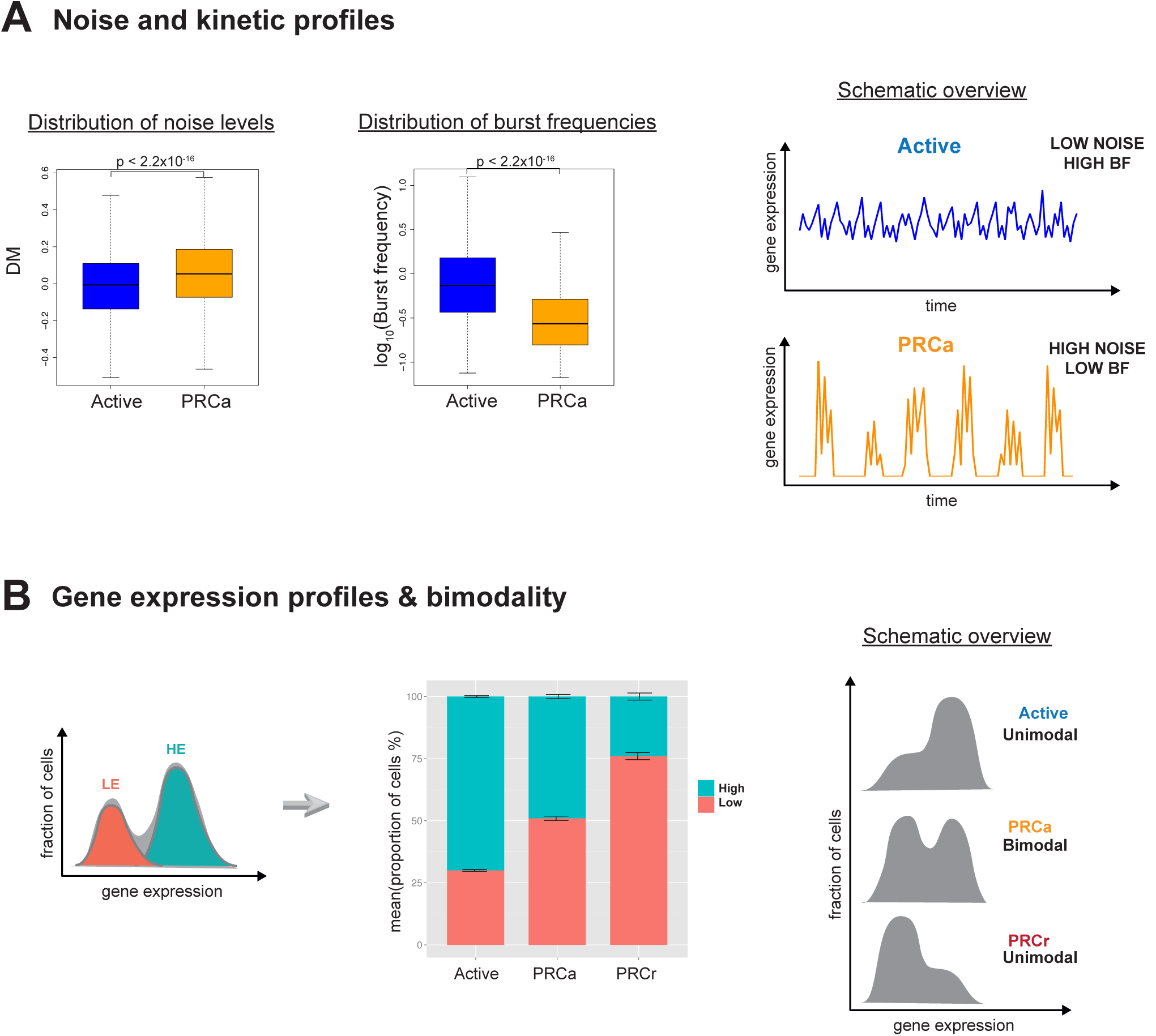
Stochastic gene expression of PRCa and active genes. (A) Comparison of PRCa and active genes reveals that PRCa genes are more variable with lower burst frequency levels than active genes (p<2.2×10^-16^ by the two-tailed Wilcoxon rank sum test). Gene expression variation is represented by DM values. (B) Expression profiles of PRCa genes show bimodal patterns. The distribution of a gene with bimodal expression is assumed to be expressed as a mixture of two normal distributions (lowly expressed (LE) and highly expressed (HE) states) (upper panel). PRCa genes have mixed cell states (on average 49% in HE and 51% in LE) indicating they are either in active state (i.e. active RNAPII and no PRC marks) or in repressed state (unproductive RNAPII and with PRC marks) consistent with cellular heterogeneity, suggested in Brookes et al.^5^. Error bars represent standard error of the mean (s.e.m).

To ensure that differences between the kinetic parameters are not driven by changes in gene expression levels between the active and PRCa groups, we extracted expression-matched genes of Active and PRCa groups (please refer to Methods). These analyses confirmed that PRCa genes have lower burst frequency and higher noise levels than Active genes (**Supplementary Fig. 3B and Supplementary Fig. 3C**). Consequently, the greater cell-to-cell variability for PRCa compared to Active genes is not driven by difference in mean expression level, but potentially linked to the presence of PRC marks themselves.

To explore whether H3K9me3 could contribute to the transcriptional heterogeneity identified at PRCa genes, we analysed H3K9me3 ChIP-Seq data of Mikkelsen et al.^17^, and found that only a few expressed PRCa genes (n=5) are marked by H3K9me3 at their promoter region (2kb centered on the TSS), making further analysis statistically impossible.

Although the literature shows that the DNA of mouse ESCs is hypomethylated, and genes that are marked by Polycomb are usually devoid of DNA methylation^18,19^, we checked the extent of DNA methylation at the PRCa gene list considered. We extracted the DNA methylation patterns at proximal promoter regions in mESCs reported in Fouse et al.^19^. Only a small proportion of genes (n=110) has DNA methylation according to this definition. Due to the small sample size, a statistical assessment will be weak, but comparison of gene expression variation profiles of these genes with the same number of PRCa genes (and same expression levels) that are unmethylated showed that the differences are not significant (Wilcoxon rank sum test *P* =0.1). This suggests no detectable effect of DNA methylation on transcriptional heterogeneity of PRCa genes (**Supplementary Fig. 3D**).

A decrease in the frequency of transcriptional bursting can manifest itself as a more bimodal pattern of gene expression across a cell population. Indeed, we observe that PRCa genes have significantly more bimodal expression profiles compared to active genes (see Methods for bimodality index calculation) (**Supplementary Fig. 3E and Figure 2B**). Assuming that the distribution of a gene with bimodal expression can be expressed as a mixture of two log-normal distributions^20^ (lowly expressed (LE) and highly expressed (HE) states), we observe that PRCa genes have mixed cell states (on average 49% of cells in HE and 51% in LE). In contrast, Active genes are mostly in the active state as expected (on average 70% in HE and 30% in LE). PRC-repressed genes with unproductive RNAPII and PRC marks, labeled as “PRCr”) are 24% in HE and 76% in LE (**Figure 2B**). Therefore, expression patterns of PRCa are in between Active and PRCr, suggesting a composite of these two states.

We should note that in our kinetic models, decay rates are set to 1 to normalize kinetic parameters so that they are independent of time^16^. To investigate whether decay rates have profound effects on kinetic parameters, we integrated published mRNA decay rates in mESCs^21^ into our kinetic model. The subtle differences in decay rates across genes did not result in major changes in the inferred kinetic parameters, leaving all major trends unaffected (**Supplementary Fig. 3F**).

### PRCa genes are important regulators in signaling pathways

To investigate potential functions of the cell-to-cell variation in gene expression in PRCa genes, we carried out KEGG pathway enrichment analysis for PRCa genes in our OS25 mESCs (see also Brookes et al.^5^). While active genes are enriched in pathways related to housekeeping functions, such as RNA transport, consistent with their uniform and stable expression across cells, PRCa genes are enriched in signaling pathways such as PI(3)K-Akt, Ras signaling and TGF-beta signaling (**Supplementary Table 1**). These signaling pathways show high levels of cell-to-cell variation compared to pathways related to housekeeping functions (**Supplementary Fig. 3G**). This may be due to transcriptomic fluctuations introduced by cytokine leukaemia inhibitory factor (LIF) signalling *via* two signaling pathways: Jak-Stat3 and PI(3)K-Akt (^22^ and **Figure 3**).

**Figure 3.**
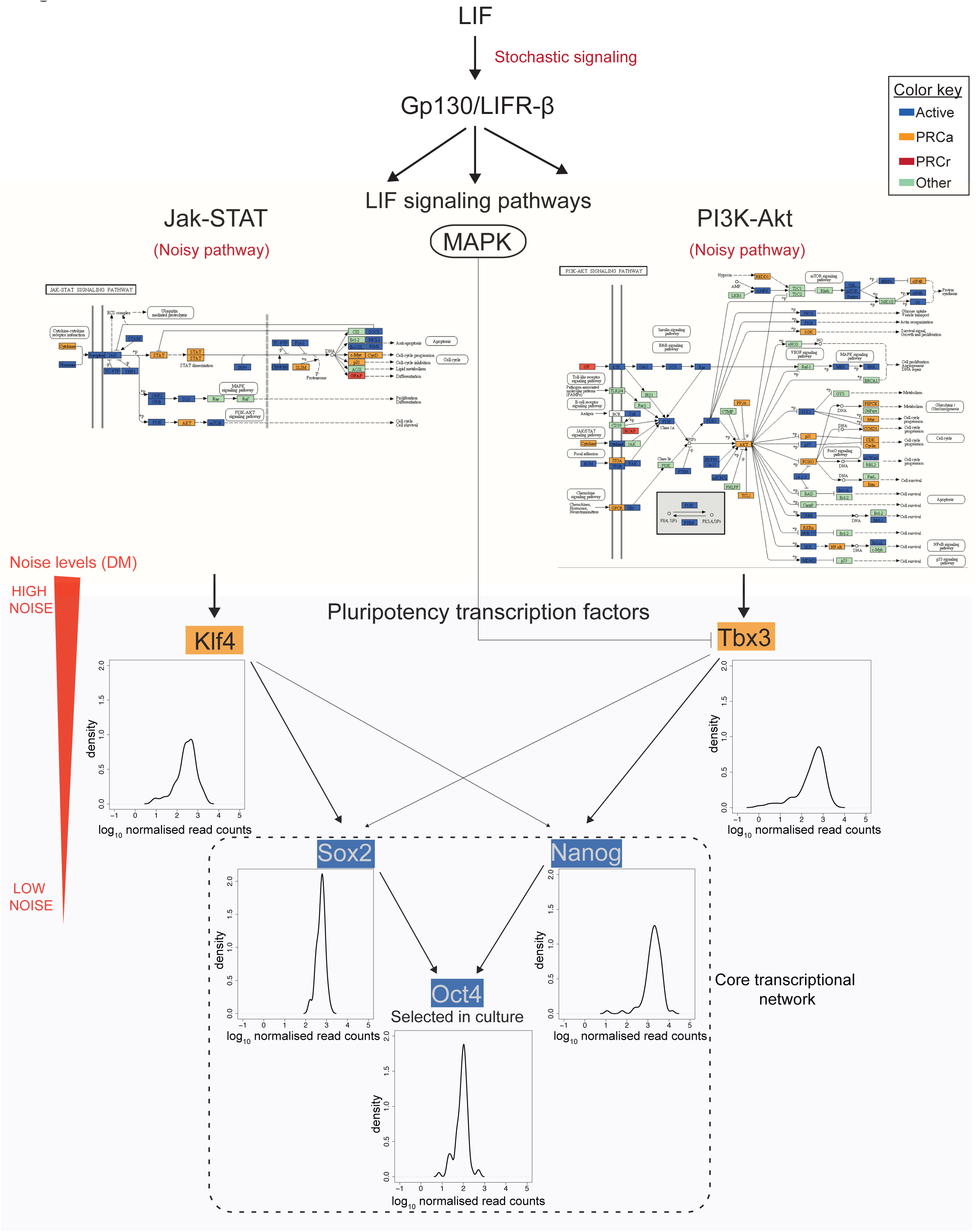
Signaling pathways that are key regulators of pluripotency in mESCs. In OS25 cells there is a selection for undifferentiated cells (high Oct4-expressing). LIF integrates signals into the core regulatory circuitry of pluripotency (Sox2, Oct4 and Nanog) via two signaling pathways; Jak-Stat and PI3K-Akt ^22^ Jak-Stat pathway activates Klf4, and PI3K-Akt pathway stimulates the transcription of Tbx3, which are both PRCa genes. The MAPK pathway antagonizes the nuclear localization of Tbx3. PRCa genes are enriched in Jak-Stat and PI3K-Akt pathways, which show high cell-to-cell variation, suggesting crucial role of PRCs in modulating fluctuations in signaling pathways that integrate LIF signals into core transcription factor network (Figure adapted from^22^).

The Jak-Stat3 pathway activates Klf4, and the PI(3)K-Akt pathway stimulates the transcription of Tbx3^22^. The expression levels of Klf4 and Tbx3, which are PRCa genes, are noisier than the pluripotency factors Nanog, Sox2 and Oct4. This pattern of noise propagation from the signaling pathways through the downstream transcriptional regulatory network is interesting, as it might indicate the role of PRCs in modulating transcriptomic fluctuations.

### Chromosomal position effects and stochastic gene expression

It is known that neighbouring genes on chromosomes exhibit significant correlations in gene expression abundance and regulation, partly due to two-dimensional chromatin domains^23-26^. Is there a similar effect of clustering by chromatin marks and noise in gene expression?

To address this, we investigated the positional effects of noise in mRNA expression using the DM values (Methods). If genes cluster together based upon their transcriptional noise, we would expect that the DM values of genes adjacent to noisy genes would be higher than those of genes adjacent to stable genes. Indeed, the noise levels of genes in the neighbourhood of noisy genes are significantly higher than those of genes that flank stable genes (*P* = 1.3×10^-4^ by the one-tailed Wilcoxon rank sum test, ±50kb of TSS, **Supplementary Fig. 4A**). This suggests that the genomic neighbourhood might influence the frequency of transcriptional bursting.

In **Figure 4A**, we show the association between chromosomal position and gene expression noise. The difference between the mean expression levels of flanking genes between noisy and stable genes is not significant (*P* = 0.7311 by the two-tailed Wilcoxon rank sum test, ±50kb of TSS), suggesting that the clusters of genes are not driven by their expression levels. The association between chromosomal position and gene expression noise was most significant at the window size of 50 kb, but weaker at a neighbourhood size of 0.5 Mb (**Figure 4A**). (Please refer to Methods for *P*-value calculation.) Thus, genes tend to be clustered into neighbourhood domains by their noise levels, ranging in size up to 0.5Mb.

**Figure 4.**
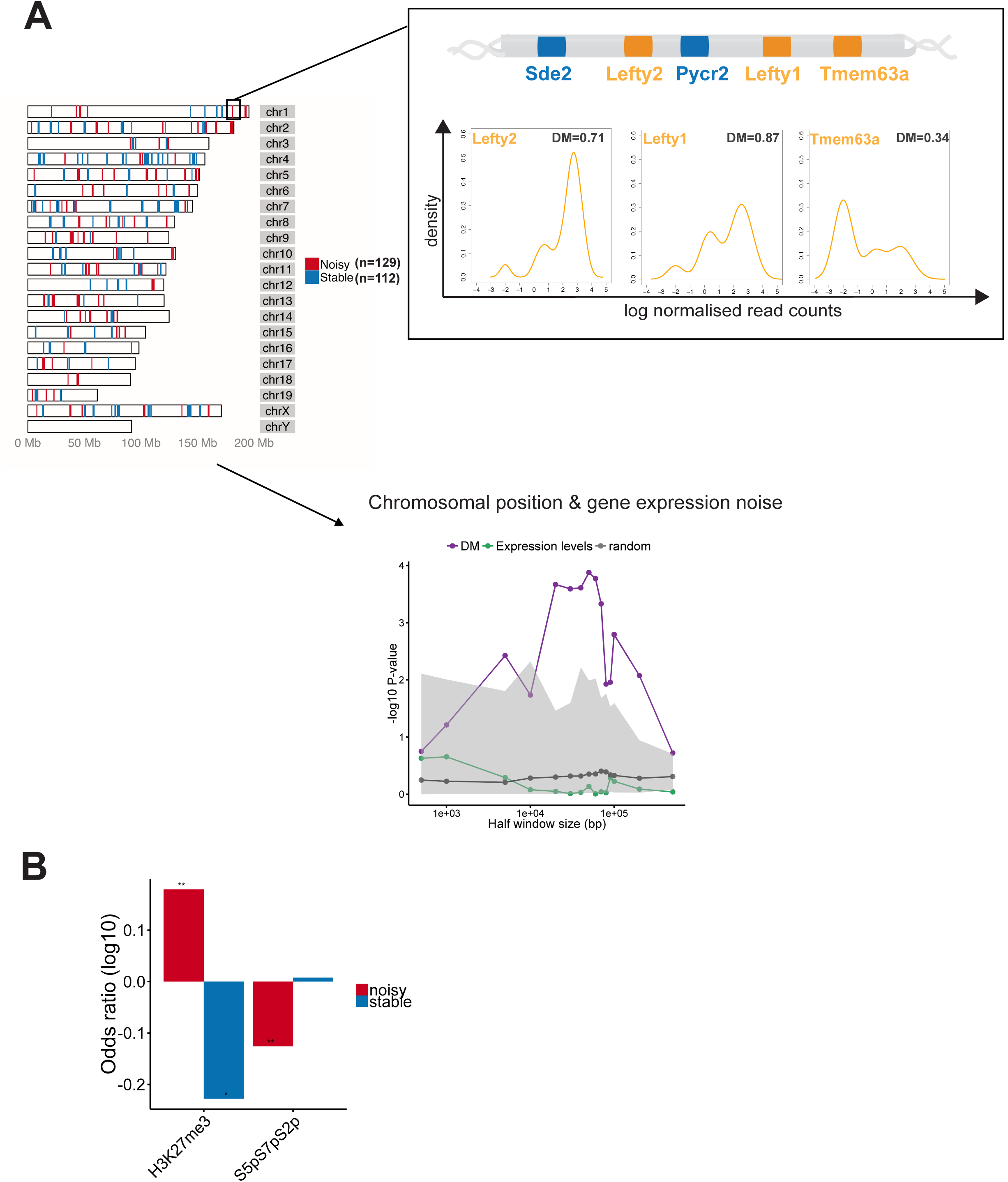
Chromosomal position effects and stochastic gene expression. (A) Maps of genes belonging to noisy and stable clusters. Chromosomal positions of genes marked by PRCs/RNAPII in the noisiest clusters. One of the noisy clusters is visualized, DM levels correlate with bimodal expression patterns. In lower panel, association between chromosomal position and gene expression noise is shown; the noise levels of genes in the neighbourhood of noisy genes are significantly higher than that of flanking genes of stable genes. As a control, we constructed 100 randomized genomes in which the positions of genes were fixed but the DM value of each gene was assigned randomly without replacement, and the same analysis was performed on each randomized genome to obtain random P-values. 2.5% quantile of random P-values, and 97.5% quantile of random P-values are shaded in gray. All data is shown on a - log(p) scale. (B) Enrichments of PRC marks/RNAPII states in noisy and stable clusters, two-tailed Fisher’s exact test; ^*^P <0.1, ^**^P < 0.05

To identify the clusters of noisy or stable genes, we performed a sliding-window analysis on the mouse genome (Methods). We found 129 noisy clusters ranging in size from 4 to 11 genes, spanning a total number of 669 genes. Similarly, 112 stable clusters (between 4 and 13 genes) with a total number of 556 genes were found (**Figure 4A**). The noise levels of genes in noisy clusters are significantly higher than that of genes in stable clusters (*P* < 2.2×10^-16^, **Supplementary Fig. 4B**) independent of mean expression levels and gene lengths (**Supplementary Fig. 4C-D)**.

Additionally, we found that DM levels correlate with bimodal expression patterns within the noisy clusters. One example is visualized in **Figure 4A**; one of the noisy clusters on chromosome 1 consists of three PRCa and two active genes. Lefty1 and Lefty2 PRCa genes, which are important in controlling the balance between self-renewal and pluripotent differentiation in mESCs, are highly variable, and also highly correlated in their gene expression. An active gene, Pycr2; Pyrroline-5-carboxylate reductase 2, is in close proximity to both Lefty1 and Lefty2, and is more variable than the Sde2 gene that lies in proximity of Lefty2 only (density profiles are shown in **Supplementary Fig. 4E**). Indeed, within the clusters, gene expression variation levels of active genes increase with the increasing number of flanking variable genes (**Supplementary Fig. 4F**). Another PRCa gene is Tmem63a, which is a transmembrane protein implicated in maintenance of pluripotency and lies near Lefty1, has high cell-to-cell variation in gene expression.

Interestingly, PRCs characterize the noisy clusters, *i.e.* PRC marks are enriched in noisy clusters rather than in stable ones. In particular, genes with H3K27me3 are enriched at noisy clusters (*P =* 1.1×10^-2^ by the two-tailed Fisher’s exact test), but depleted at stable clusters (*P* = 5.9×10^-2^, **Figure 4B**). Since PRCs are tightly associated with RNAPII states, we examined differences between the RNAPII state of genes between noisy and stable clusters. We found that genes marked by active elongating RNAPII (S5pS7pS2p) are depleted at noisy clusters (*P* = 1.3×10^-3^ by the two-tailed Fisher’s exact test, **Figure 4B**), supporting the view that elongating RNAPII modifications promote stable gene expression. Together, noisy clusters are characterized by the presence of PRC marks and the absence of active elongating RNAPII, while stable clusters are characterized by the absence of PRCs.

### Gene and enhancer clustering in 2D and 3D

Next, we analyzed whether PRCa genes are proximal to fully repressed Polycomb genes, which could eventually increase their sensitivity to Polycomb repression. Linear spatial proximity between PRCa genes and PRCr genes is significantly closer than the median distance between randomly chosen genes and PRCr genes (*P* = 2×10^-2^, **Figure 5A**) (Methods). Interestingly, PRCa genes are also in close proximity to Active genes (*P* < 0.0001, **Supplementary Fig. 4G**), while Active genes are distal from PRCr genes (*P* = 5×10^-3^, **Supplementary Fig. 4H**), suggesting a 2D spatial arrangement of these genes as Active-PRCa-PRCr (as visualized in **Figure 5A**).

**Figure 5.**
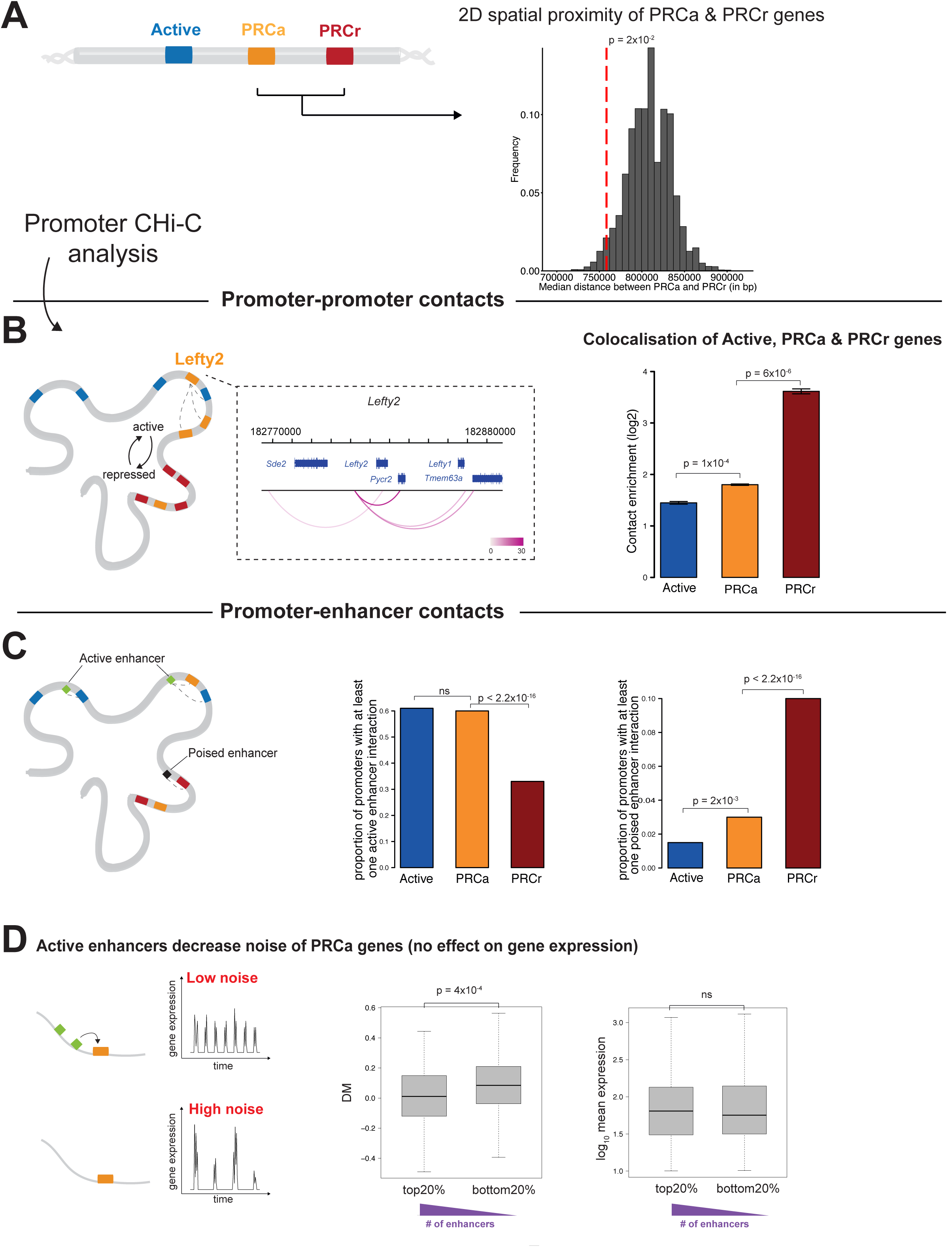
Effects of 2D and 3D neighborhood on transcriptional kinetics (A) Histogram of simulated median distances under a null model assuming no positional preference in the neighbourhood of PRCr genes. The observed median distance of PRCa genes to their nearest neighbor in the PRCr group, depicted by vertical dashed red line, are significantly less than expected by chance (*P* = 2×10^-2^). (B) Analyzing mESC Promoter Capture Hi-C data reveals that PRCa genes have a strong enrichment for long-range contacts between promoters with levels in between PRCr and active genes. Error bars represent s.e.m. (C) PRCa genes have significantly more interactions with active enhancers compared to PRCr genes (P < 2.2×10^-16^). In contrast, interactions with poised enhancers are mainly observed for PRCr genes rather than PRCa (P < 2.2×10^-16^). (D) Occurrence of interactions with active enhancers decrease noise of PRCa genes independent of mean expression levels.

We next asked whether the linear genomic position effects of PRCs are reflected in the 3D genome organization in ESCs. Recently, Schoenfelder et al.^27^ found that PRC1 acts as a major regulator of ESC genome architecture by organizing genes into three-dimensional interaction networks. They generated mouse ESC Promoter Capture Hi-C (CHi-C) data^28^, and analysed it using the GOTHiC (Genome Organization Through Hi-C) Bioconductor package. This yielded a strong enrichment for long-range contacts between promoters bound by PRCs.

We applied the same approach to this dataset using our gene list. We found that there is a strong enrichment for long-range promoter-promoter contacts for both PRCa and PRCr genes (**Figure 5B**). Interestingly, PRCr genes have significantly stronger contact enrichment than PRCa genes in mESCs (one-tailed t-test *P* = 6.3×10^-6^). PRCa genes are in between PRCr and Active genes; they have stronger contact enrichment than Active genes (one-tailed t-test *P* = 1×10^-4^) (**Figure 5B**).

In **Figure 5B**, the promoter contacts of the aforementioned noisy cluster PRCa gene Lefty2 is visualized. It is in contact with the other PRCa genes Lefty1 and Tmem63a, and it has strong connectivity with the active Pycr2 genes. These contacts may affect Pycr2’s frequency of transcriptional bursting, and thereby tune expression noise.

In terms of the promoter preferences of gene sets, it is interesting to note that PRCa promoters interact equally with promoters of PRCr, PRCa and Active genes (**Supplementary Fig. 4F**). However, PRCr promoters have a distinct preference for other PRCr promoters (two-tailed Fisher’s exact test P < 2.2×10^-16^).

We next investigated contacts between PRC promoter classes with putative regulatory (non-promoter) elements; enhancers that are described as in Schoenfelder et al. ^27^; active (H3K4me1 and H3K27ac), intermediate (H3K4me1) or poised (H3K4me1 and H3K27me3) enhancers. We found that PRCa genes have significantly more interactions with active enhancers compared to PRCr genes (P < 2.2×10^-16^) (**Figure 5C**). In contrast, interactions with poised enhancers are mainly observed for PRCr genes rather than PRCa (P < 2.2×10^-16^).

Further, we asked whether interactions with enhancers affect transcriptional profiles of PRCa genes at the single cell level. Interestingly, we found that interactions with active enhancers decrease noise in gene expression of PRCa genes. Sorting the PRCa genes based on the number of active enhancer interactions shows that more interactions lead to less noise in gene expression (two-sided Wilcoxon test *P* = 4×10^-4^). This stabilization of expression through active enhancers is independent of mean expression levels (**Figure 5D**).

In summary, these findings show that 3D genome architecture correlates with chromatin state, and may influence noise in gene expression. This holds both in terms of promoter-promoter and enhancer-promoter interactions.

### Ring1A/B double knockout affects transcriptional profiles of PRC-bound genes

To test whether noise in gene expression can be linked to Polycomb regulation mechanistically, we utilized conditional Ring1B double knockout (in Ring1A-/- background) mES cells. These cells lack Ring1A, and have a tamoxifen-inducible conditional Ring1B deletion (**Supplementary Fig. 5A** and Methods). We confirmed Ring1B deletion 48 hours post-tamoxifen induction, and generated single cell RNA-sequencing data for both untreated (Ring1A single KO) and tamoxifen-treated double KO (Ring1A and Ring1B dKO) mES cells (see Methods). In these conditions, Ring1B protein is lost ~48h, and H2Aub1 modification is no longer detected on chromatin and Polycomb repressed genes are derepressed without loss of pluripotency factors Nanog, Oct4 and Rex1^5,8,29^.

We compared the changes in mean expression at PRCr, PRCa and active genes. We found that PRCr show substantial derepression after Ring1A/B dKO (**Figure 6**), as expected from bulk mRNA-seq/microarray data^5,29^. The mean expression change at PRCa genes is lower than at PRCr (*P =* 4.1x10^-9^ by the two-tailed Wilcoxon rank sum test) (**Figure 6**) more likely due to the fact that they are already expressed to some extent in untreated cells. Nevertheless, changes in mean expression at PRCa genes are higher than at active genes (P = 2×10^-7^) (**Figure 6** and **Supplementary Fig. 5B**). Increased expression of PRCa genes upon Ring1A/B dKO recapitulates previous findings using bulk transcriptomic analyses^5,8^.

**Figure 6.**
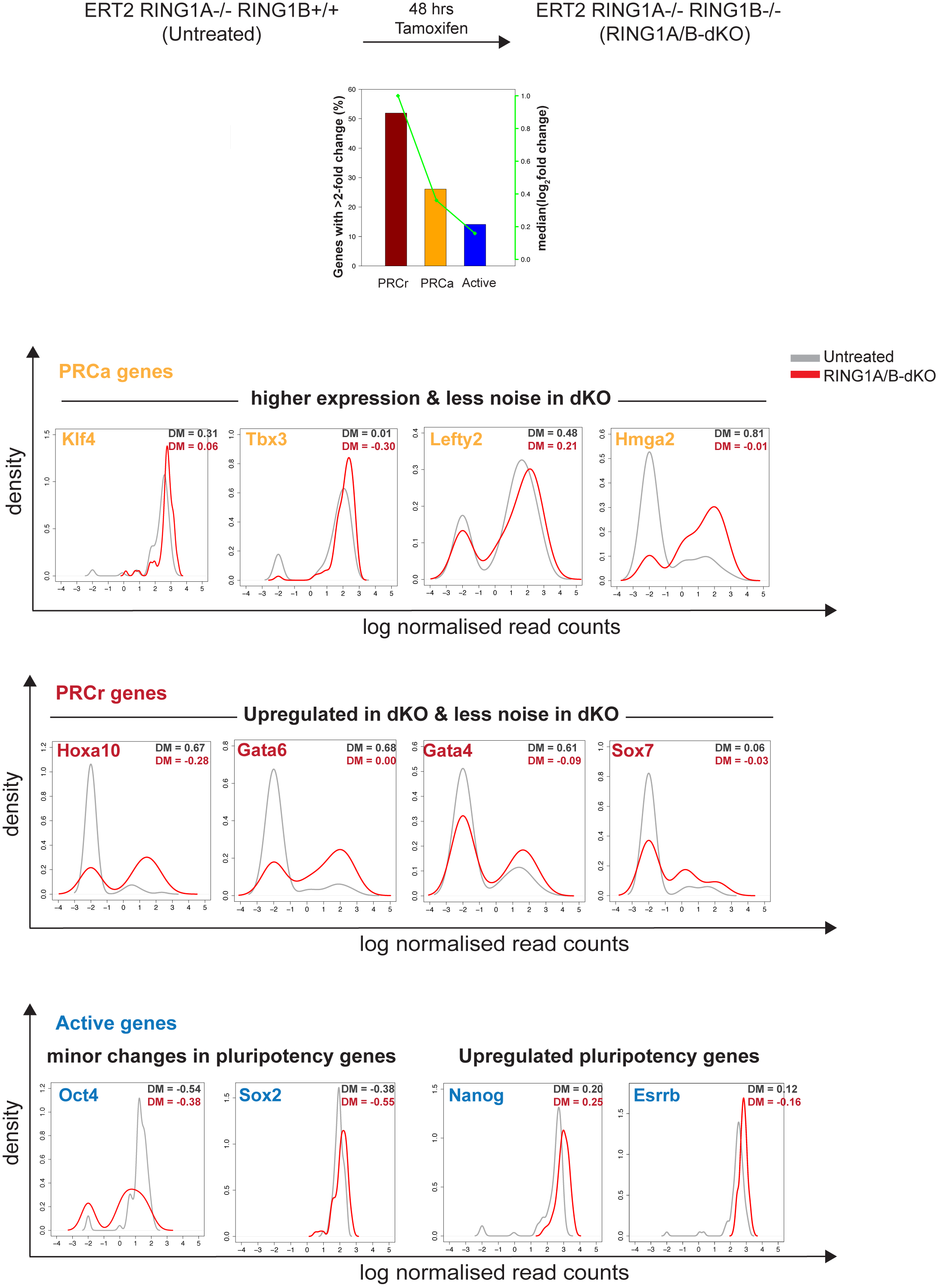
Single cell profiling of Ring1A/B double knockout mES cells. PRCr show substantial derepression after Ring1A/B dKO. The mean expression change at PRCa genes is lower than at PRCr, in contrast, is higher than at active genes. Comparison of noise levels show that there is a more pronounced decrease in noise levels at PRCa genes compared to active genes. Gene expression profiles of some important genes for ESC biology are shown. Key pluripotency transcription factors Klf4 and Tbx3 are more expressed and less noisy in Ring1A/B dKO cells. Other transcriptional regulators such as Hmga2 and Hdac2 become upregulated after dKO. Consistently, key differentiation markers such as Gata4, Gata6 are upregulated. Among active pluripotency factors, Oct4 and Sox2 show minor changes in expression (mean expression levels are not significantly different), and Nanog and Esrrb are upregulated.

Importantly, comparison of noise levels shows that there is a more pronounced decrease in noise levels at PRCa genes compared to active genes upon Ring1A/B dKO (*P* = 4×10^-3^) (Supplementary Fig. 5C). This supports our findings that Polycomb tunes gene expression noise. Additionally, there is a more pronounced decrease in bimodality at PRCa genes (**Supplementary Fig. 5D**), while burst frequency levels decrease more significantly at active genes (**Supplementary Fig. 5E**).

Among PRCa genes, key pluripotency transcription factors Klf4 and Tbx3 and other transcriptional regulators (such as Hmga2 and Hdac2) important for ESC biology become upregulated and show less noisy profiles after Ring1A/B dKO (gene expression profiles are shown in **Figure 6**). Additionally, key differentiation markers such as Gata4, Gata6, which are PRCr genes, are upregulated upon dKO (**Figure 6**), implying that a Polycomb KO could make cells more prone to differentiation (as expected from^5, 29^). The same pattern of differential gene expression is also observed in the bulk RNA-sequencing data. Taken together, these findings indicate the key role of Polycomb in regulating transcriptional profiles of PRC-bound genes.

We observe that non-PRC targets (i.e. active genes) show subtle trends in change in gene expression; expression levels of active pluripotency factors such as Oct4 and Sox2 show minor changes in gene expression. In contrast, Nanog and Esrrb are upregulated (**Figure 6**), suggesting that Polycomb may indirectly control the expression of genes specifically associated with pluripotency. Expression patterns of all these genes can be found at http://www.ebi.ac.uk/teichmann-srv/espresso/.

## Discussion

It is well understood how post-translational modifications of histones, including acetylation, methylation, phosphorylation and ubiquitination, modulate chromatin structure, thereby affecting the regulation of gene expression levels^30^. It is much less well understood how chromatin status is related to the kinetics of transcription in terms of transcriptional bursting. Differences in stochastic gene expression lead to different degrees of cell-to-cell variation in expression levels, even for genes with the same mean expression across an ensemble of cells. Recent molecular studies have shown that individual cells can show substantial differences in both gene expression and phenotypic output^31,32^. Genetically identical cells may still behave differently under identical conditions^33^. This non-genetic variability is mainly due to cell-to-cell variation in gene expression^34,35^, which relates to each gene’s chromatin status ^36^. Noisy or stochastic gene expression profiles may play an important role in the regulation of ES cells^37^.

In this work, we focus on histone modifications that are mediated by Polycomb repressive complexes and investigate their relationship with stochastic gene expression in mES cells. Earlier work indicated that expression of Polycomb target genes negatively correlates with levels of H3K27me3, and suggested that dynamic fluctuations in chromatin state are associated with expression of certain Polycomb targets in pluripotent stem cells^38^. Although PRCs are known to exert a repressive effect, interestingly, the cohort of PRC-bound genes contains not only silent genes, but also genes with intermediate and high expression^5^. A large range of expression levels at PRC-target genes is observed in published mRNA data sets^5,39^ and substantial expression has been previously observed at PRC2-target genes^40,41^. The moderate to high expression levels at some PRC-bound genes allow us to reliably quantify gene expression variation (which is not possible if expression is too low).

Here, benefiting from the power of single cell RNA-seq analysis, we show that PRCa genes have greater cell-to-cell variation in expression than their non-PRC counterparts, suggesting that they switch between on and off states in a more dramatic way. Along the same lines, their expression patterns are more likely to be bimodal, suggesting a composite of active and PRC-repressed states at the single cell level. These findings indicate the role of Polycomb in modulating frequency of transcriptional bursting and thereby tuning gene expression noise.

Transcriptional bursts that arise from random fluctuations between open and closed chromatin states of a gene are one of the major sources of gene expression noise in eukaryotes ^42^ Since these fluctuations are modulated by transcription factors, nucleosomes and chromatin remodelling enzymes, we can speculate that gene expression noise may be linked to chromosomal position through shared chromatin domains with specific characteristics such as histone modifications. Consistent with this notion, several studies using a reporter transgene integrated in multiple loci have shown that gene expression noise varies with chromosomal position in yeast and mammalian cells^43-48^.

However, large-scale studies measuring noise in protein expression of endogenous genes could not find a strong association between chromosomal position and gene expression noise in yeast^34,49^. This discrepancy might be due to gene-specific confounding factors and different statistics to examine the association. For example, essential genes with low noise derived from the same datasets of the large-scale proteomic studies are clustered into neighbourhood domains with low nucleosome occupancy^23^. More importantly, noise in protein expression is not a good measure for examining the effect of chromatin regulation on transcriptional bursting since slowly degrading proteins can buffer transcriptional noise at the protein levels^46^. Given the lack of high-throughput measurements of noise in mRNA expression of endogenous genes in eukaryotes, it is not clear if genes are distributed across the genome by their noise levels and which chromatin features modulate the chromosomal position effects.

Analysis of the chromosomal position effects of noise reveals that genes are significantly clustered according to their noise levels, which are mainly modulated by PRCs. Interestingly, across the chromosomes, we found that PRCa genes are in close proximity to fully repressed PRC-targets. This could increase their sensitivity to PRC-repression, and explain their ability to switch between active and repressed states in a more dramatic way than other genes.

In addition to 2D spatial proximity of genes, long-range regulatory interactions have a key role in gene expression control^50^. Recently, analyzing mouse ESC Promoter Capture Hi-C (CHi-C) data^28^, Schoenfelder et al. showed that PRC1 acts as a major regulator of ESC 3D genome architecture^27^. Applying the same methodology, we show that there is a strong enrichment for long-range promoter-promoter contacts for both PRCa and PRCr genes. Interestingly, interactions with active enhancers decrease gene expression noise (but not mean expression levels) of PRCa genes, suggesting that 3D genome architecture has a key role in controlling gene expression noise.

To further decipher the role of PRCs in regulating gene expression and noise, we performed single cell RNA-Sequencing for both PRC-expression (Ring1A-KO, untreated) and PRC-deleted (Ring1A/B-dKO, tamoxifen-treated) mESCs. We observe substantial derepression of PRC-bound genes after Ring1A/B-dKO as expected. Mean expression changes at PRCa genes are significantly lower than at PRCr genes, supporting the fact that they are already expressed in untreated ES cells. Moreover, in terms of noise profiles, we observe a significant decrease in noise levels of PRCa genes compared to Active genes. This genetic validation supports our findings that polycomb plays a key role in modulating the kinetics of stochastic gene expression.

## Methods

### Single-cell RNA-sequencing of mouse OS25 ES cells

Mouse ES-OS25 cells were cultured as described before^5^. For single cell sequencing libraries were prepared according to Fluidigm manual “Using the C1 Single-Cell Auto Prep System to Generate mRNA from Single Cells and Libraries for Sequencing“. OS25 cell suspension was loaded on 10-17 micron C1 Single-Cell Auto Prep IFC, Fluidigm, cDNA was synthesized in the chip using Clontech SMARTer kit and Illumina sequencing libraries were prepared with Nextera XT kit and Nextera Index Kit (Illumina). Libraries from 96 cells were pooled and sequenced on 4 lanes on Illumina HiSeq2000 using 100bp paired-end protocol.

### Mapping reads

For each cell, paired-end reads were mapped to the *Mus musculus* genome (GRCm38) using GSNAP with default parameters^51^. Next, uniquely mapped reads to the genome were counted using htseq-count (http://www-huber.embl.de/users/anders/HTSeq/) and normalized with size factors using DESeq^52^.

### Classification of genes based on ChIP-Seq profiles

To integrate ChIP-Seq data with single-cell RNA-Seq, we mapped 18,860 UCSC known gene IDs from Brookes et al.^5^ to Ensembl IDs using BioMart^53^. Then, we categorized the genes based on Brookes et al. classification: (1) “Active” genes (n=4,732) are defined as those without PRC marks (H3K27me3 or H2Aub1) but with active RNAPII (S5pS7pS2p), (2) “PRCa” (n=1,263) genes are marked by PRCs (H3K27me3 or H3K27me3 plus H2Aub1) and active RNAPII, (3) “PRCr” genes (n=954) have both PRC marks (H3K27me3 and H2Aub1), unproductive RNAPII (S5p only and not recognized by antibody 8WG16) and not expressed in Brookes et al.’s bulk mRNA data (bulk mRNA FPKM<1). We should note that vast majority of PRCa and PRCr genes are H3K4me3 positive (1248/1263 PRCa, and 938/954 PRCr) (see Brookes et al.^5^)

We focus on Active and PRCa genes with moderate to high mRNA abundance and therefore we remove genes that have mean normalized counts lower than 10. Thus, in the final gene set, there are 4,483 Active genes and 945 PRCa genes.

For H3K9me3, reads from Mikkelsen et al.^17^ were mapped to mouse genome (mm9, July 2007) using Bowtie2 v2.0.5^54^, with default parameters. Enriched regions were identified with BCP v1.1^55^ in Histone Mark mode, using as control H3 from Mikkelsen et al., processed in the same way. Genes were defined as positive for H3K9me3 at their promoter or gene body when an enriched region was overlapping with a 2kb window around the TSS or between the TSS and TES, respectively.

### Inference of transcriptional kinetic parameters via modeling single-cell RNA-seq data

To explore kinetics of stochastic gene expression, we fitted a Poisson-beta model as described previously^16^. Poisson-beta model is an efficient way to describe the long-tailed behavior of mRNA distribution resulting from occasional transcriptional bursts as well as to explain expression bimodality of genes with low burst frequency. Transcriptional kinetic parameters are characterized by two parameters, burst size is described as the average number of synthesized mRNA molecules while a gene remains in an active state and burst frequency is the frequency at which bursts occur per unit time. To ensure that the parameters are statistically identifiable, a goodness-of-fit statistic is applied as described in^16^. Out of 5,428 genes (active and PRCa), 4,526 genes (83%) have identifiable estimates of kinetic parameters. We focus henceforth on these genes in analysis of burst size and frequency.

We should note that our kinetic analyses do not account for technical noise as our data do not contain external spike-in molecules (the only way to incorporate technical noise in our kinetic model). Therefore, we addressed this point by focusing on moderately or highly expressed genes with an expression cutoff of 10. The assumption is that technical noise for these genes is small enough to estimate kinetic parameters accurately. We should also note that our results are robust to changes in selection of expression cutoff (**Supplementary Fig. 3A**).

### Controlling for expression levels in kinetic models

To control for expression levels for PRCa and Active gene sets, we extracted expression-matched sets of active and PRCa genes using “matching” function in R “arm” package with default settings. In this way, an active gene is matched to a PRCa gene that has the closest mean expression level.

### Calculation of DM as a measure of gene expression variability

Widely used measures for quantifying gene expression variation in mRNA expression such as the CV and Fano factor are not suitable for assessing differences in gene expression variation between genes because they depend strongly on gene expression levels and gene length. To properly account for the confounding effects of expression level and gene length on variation, we first computed a mean corrected residual of variation by calculating the difference between the observed squared CV (log10 transformed) of a gene and its expected squared CV. As a second step to correct for the effect of gene length on the mean corrected residual of variation, we calculated the difference between the mean corrected residual of the gene and its expected residual, which is referred to as DM^11^. The expected squared CV or the expected residual was approximated by using a running median.

### Calculation of bimodality index

Bimodality index was calculated as described previously by Wang et al.^20^. The distribution of a gene with bimodal expression is assumed to be described as a mixture of two normal distributions with equal standard deviation. Proportions of observations in two components were estimated using R package ‘mclust’.

### Identifying noisy and stable genes across mouse chromosomes using DM values

To investigate the position effects of noise in mRNA expression using DM values, we first sorted all expressed genes (n=11,861) in descending order according to their DM values and chose the top 20% as “noisy” genes and the bottom 20% as “stable” genes. For each gene, we counted the number of noisy (or stable) genes (excluding the focal gene) in the neighbourhood of the gene (±0.5kb ~ 500bp of the transcription start site (TSS) of the focal gene).

While investigating the association between chromosomal position and gene expression noise, as a control, we constructed 100 randomized genomes in which the positions of genes were fixed but the DM value of each gene was assigned randomly without replacement, and the same analysis was performed on each randomized genome. The *P* values observed in the real genome are less than the median of *P* values found in the randomized genomes at all neighbourhood sizes and even less than the 2.5% quantile of random *P* values at the neighbourhood sizes between 20kb and 0.2Mb (**Figure 4A**).

### Identifying clusters of genes by a sliding-window approach

To identify the clusters of noisy or stable genes in the mouse genome, we used a sliding-window approach^56^ with a window size of four genes. Given a set of genes having valid DM values, a window starts from the first gene of each chromosome and keeps shifting right by one gene until it reaches the end of the chromosome. We ignored windows having a distance between TSSs of the first and fourth gene of the windows larger than (window size – 1) × 0.5Mb. We measured the overall noise of each window by summing rolling means of the DM values of two consecutive genes within the window. We then calculated this noise score of randomly chosen four genes, and repeated this process 100,000 times, yielding a null distribution of the overall noise score of a window. We called a window to be significantly noisy (or stable) if its noise score is above 97.5% of randomized windows (or below 2.5% of randomized windows). Finally, we merged all overlapped noisy (or stable) windows to construct a set of noisy (or stable) clusters.

The total number of genes in noisy clusters found in the mouse genome is not significantly higher than that of 1,000 randomized genomes (empirical *P* = 0.3996). In contrast, the total number of genes in stable clusters is significantly lower than expected by chance (empirical *P =* 1.0×10^-3^), suggesting that the stable clusters are relatively rare.

### Testing the spatial proximity between PRCa and PRCr genes

To test whether PRCa genes are in the neighbourhood of PRCr genes, we calculated the distance for each gene in the PRCa group (1,263 genes) to its nearest neighbour in the PRCr group (954 genes) using TSSs. The observed mean and median distance were tested against a null model assuming no positional preference of PRCa genes in the neighbourhood of PRCr genes. We observed that a majority of genes not expressed in mESCs are distal from Active/PRCa/PRCr genes. To correct for the effect of these inactive genes, we defined a background set of genes as ones belonging to Active, PRCa, or PRCr genes. We randomly sampled 1,263 genes from the background set by excluding genes that are in the PRCr group or in the chromosomes on which the 954 PRCr genes are not located, and calculated the mean and median distance between the randomly chosen genes and PRCr genes. We repeated this process 10,000 times and computed the empirical *P*-values of the observed mean and median distance based on a null distribution of simulated distances.

### Promoter-promoter contacts and contact enrichment analysis

Significant promoter-promoter and promoter-genome interactions in WT ESC were obtained from Schoenfelder et al.^27^. Short range intra-chromosomal contacts were excluded by filtering contacts separated by <10 Mb. To measure the enrichment of contacts within a set of promoters, 100 random promoter sets were generated with comparable pair-wise distance distributions to the experimental set. Contact enrichment was derived by dividing the number of contacts in the experimental set by the average number of contacts in the control sets. For each experimental set, we calculated the contact enrichment using three independent control sets and showed the mean contact enrichment and the standard deviation. Contact enrichment differences were evaluated using one-tailed t-tests.

### Gene Ontology and KEGG pathway analyses

Annotation of KEGG pathways^57^ and their associated genes were retrieved using Bioconductor Package KEGGREST. Enrichment of KEGG pathways was assessed by Fisher’s exact test in R Stats package and P-values were adjusted for multiple testing by calculating false discovery rates.

### Ring1A/B double knockout cells and mRNA sequencing

Ring1A/B double knockout cells^29^ (kind gift from Neil Brockdorff, which have been authenticated before) with constitutive Ring1A knockout and tamoxifen-inducible conditional Ring1B knockout were cultured on mitomycin-inactivated feeders in DMEM (lacking pyruvate; Gibco), supplemented with 10% batch-tested FCS (Labtech), 50mM β-mercaptoethanol, L-glutamine (Gibco), Sodium Pyruvate (Gibco), Non-essential amino acids (Gibco), Penicillin/Streptomycin (Gibco) supplemented with 1000U/ml LIF (Milipore)^29^. These cell lines have been tested and were found to have no mycoplasma contamination. Feeders in Ring1AB dKO mES cells (untreated and tamoxifen-treated) are depleted using Feeder removal MicroBeads (Miltenyi Biotec). To induce Ring1b knockout, cells are cultured in media containing 800nM 4-hydroxytamoxifen (Sigma) for 48 hours and confirmed using genomic DNA isolation and PCR across Cre-excised region^8,29^. Primer information^29^ is listed below.

Ring1b-s3 AAGCCAAAATTTAAAAGCACTGT

Ring1b-4681 ATGGTCAAGCAAACATGAAGGT

Ring1b-as4 TGAAAAGGAAATGCAATGGTAT

All cells are processed on C1 Single Cell Auto Prep System (Fluidigm; 100-7000 and 100-6209) using medium sized C1 mRNA-Seq chips (10-17 μm; 100-5670) with ERCC spike-ins (Ambion; AM1780) following the manufacturers protocol (100-5950 B1) requiring SMARTer kit for Illumina Sequencing (Clonetech; 634936). Single cell libraries were made using Illumina Nextera XT DNA sample preparation kit (Illumina; FC-131-1096) after cleanup and pooling using AMPure XP beads (Agencourt Biosciences; A63880). Each library is sequenced on single HiSeq2000 lane (Illumina) using 100bp paired-end sequencing.

We also generated standard bulk RNA-sequencing for each condition. Bulk RNA-sequencing libraries were prepared and sequenced using the Wellcome Trust Sanger Institute sample preparation pipeline with Illumina’s TruSeq RNA Sample Preparation v2 Kit as described before^11^. We observed that average single cell expression levels recapitulated the bulk gene expression levels with a Spearman rank correlation coefficient of 0.89 and 0.88 for untreated and dKO conditions respectively.

## URLs

GOTHiC Bioconductor package,
http://www.bioconductor.org/packages/release/bioc/html/GOTHiC.html

## Acknowledgements

We are grateful to Neil Brockdorff for the kind gift of Ring1A/B dKO cells. We acknowledge funding from the Wellcome Trust Strategic Award on Mouse Gastrulation, the Lister Institute and the Helmholtz Foundation.

## SUPPLEMENTARY FIGURE LEGENDS

**Figure S1.** (A,B) Single-cell data statistics: over 80% of reads were mapped to the *Mus musculus* genome (GRCm38) and over 60% to exons. (C) Quality control analysis for single cells. (D) Heatmap showing expression profiles of key pluripotency factors and differentiation markers in OS25 cells. There is homogenously high expression of pluripotency genes, and all differentiation markers are consistently “off”. This indicates that OS25 cells are all in a pluripotent state. (E) OS25 cells are shown together with other mESCs cultured in serum+lif and 2i from Kolodziejczyk et al. 2015. OS25 cells are more similar to the subpopulation of pluripotent serum cells, rather than the subpopulation of serum cells that are either “primed for differentiation” or “on the differentiation path”. (F) Average single-cell expression is highly correlated with bulk RNA-Seq (data from Brookes et al.^5^), Spearman’s correlation coefficient is 0.87.

**Figure S2.** (A) Squared coefficient of variation (CV^2^) vs. average normalized read count of genes are shown (x and y-axes log10-scale). As gene expression levels increase, genes are more likely to show lower levels of variation. Variable genes are in red color and cell cycle genes (from Gene Ontology and Cycle base database) are in green color. (B) Gene expression variation components are unraveled by applying a recent method ^13^, which uses Gaussian Process Latent variable models in single-cells (scLVM). It is a two-step approach that first reconstructs cell cycle state and then uses this information to obtain “corrected” gene expression levels. Cell cycle contribution to variation is around 1% on average. In the lower panel, gene expression profiles for Aurka, a cell cycle gene and Klf4, a pluripotency transcription factor, are shown. After cell cycle regression, profile of Aurka becomes more homogeneous, whereas Klf4 remains uncorrected. (C) Clustering of 90 cells based on cell cycle G2/M stage markers: there are two groups: one with high expression of G2 and M genes and the other with low expression of these genes. (D) Clustering after cell cycle correction: cell cycle effect is removed leading to more homogeneous expression distribution of these genes across the cells.

**Figure S3. (A)** Distribution of DM and burst frequency levels across different cutoffs of gene expression. Two-sided Wilcoxon rank-sum test P-values for differences of DM between Active and PRCa genes are: P<2.2×10^-16^, P<2.2×10^-16^, P<2.2×10^-16^, P=6.7×10^-15^, P=2.3×10^-13^, P=1.8×10^-13^, P=8.1×10^-12^, P=1.7×10^-13^, P=2.2×10^-12^ and P=5.6×10^-11^ for gene expression cutoffs 10, 20, 30, 40, 50, 60, 70, 80, 90 and 100, respectively. For burst frequency levels, all P-values are P<2.2×10^-16^. (B) Expression matched sets of Active and PRCa genes show that differences in DM and burst frequency levels are independent of gene expression levels. (C)–log10 P-values are shown for differences between DM levels and BF levels of expression-matched Active and PRCa groups across different expression cutoffs. The number of PRCa genes (that are expression-matched to same number of Active genes) are 666, 603, 540, 479, 427, 374, 341, 304, 280 and 262 for expression cutoffs of 10, 20, 30, 40, 50, 60, 70, 80, 90 and 100, respectively. (D) Comparison of gene expression variation profiles of methylated and unmethylated PRCa genes suggests that DNA methylation has no pronounced effect on transcriptional heterogeneity in mESCs. (E) PRCa genes are more bimodal than active genes. (F) Taking into account mESC degradation rates and including them into our kinetic models does not result in major changes in kinetic parameters, thereby yields similar findings. (G) Median DM of KEGG signaling pathways PI(3)K-Akt, Ras signaling and TGF-beta signaling (shown in purple color) are significantly higher compared to median DM levels of pathways related to housekeeping functions, such as RNA transport and Ribosome (shown in green).

**Figure S4.** (A) Noise levels of genes in the neighborhood of noisy genes are significantly higher than those of genes that flank stable genes. (B) The difference of DM between noisy and stable genes is significant (P < 2.2×10^-16^). (C) The difference of gene length between noisy and stable genes is not significant (*P* = 0.1563). (D) The difference of mean expression levels between noisy and stable genes is not significant (*P* = 0.8485 by the two-tailed Wilcoxon rank sum test). (E) Gene expression profiles and DM levels of active genes; Sde2 and Pycr2 in one of the noisy clusters are shown. (F) Noise levels of active genes flanked by zero, one and two variable genes: genes flanked by two variable genes show highest levels of variation, while genes flanked by zero variable genes are more stable than other groups. (G) The observed median distance of Active genes to their nearest neighbor in the PRCa group, depicted by vertical dashed red line, are significantly less than expected by chance (P < 0.0001). (H) The observed median distance of Active genes to their nearest neighbor in the PRCr group, depicted by vertical dashed red line, are significantly higher than expected by chance (*P* = 5×10^-3^). (I) Promoter preferences of gene sets: PRCr promoter preferences are different; PRCr genes are more likely to interact with PRCr promoters than PRCa genes (two-tailed Fisher’s exact test P < 2.2×10^-16^). Similarly, PRCa are more likely to interact with PRCr promoters than Active genes (two-tailed Fisher’s exact test *P* = 1×10^-3^)

**Figure S5.** (A) Schematic layout of Ring1B locus (UCSC mm10 reference assembly) and PCR primers to confirm Ring1b knockout. PCR amplification of genomic DNA from untreated (Ring1AKO) and tamoxifen-treated (Ring1ABdKO) to confirm Ring1B knockout are shown. Expected fragment size in untreated and Tamoxifen treated samples listed on right. (B) PRCa genes have a more pronounced change in mean gene expression and (C) noise levels (D) bimodality patterns than active genes. (E) Decrease in burst frequencies are more pronounced at active genes.

**Table S1.** KEGG pathway enrichment of PRCa genes.

| Pathways | FDR |
| --- | --- |
| path:mmu04014 Ras signaling pathway | 0.0002 |
| path:mmu04060 Cytokine-cytokine receptor interaction | 0.0010 |
| path:mmu04151 PI3K-Akt signaling pathway | 0.0023 |
| path:mmu05206 MicroRNAs in cancer | 0.0023 |
| path:mmu05200 Pathways in cancer | 0.0036 |
| path:mmu04015 Rap1 signaling pathway | 0.0043 |
| path:mmu00604 Glycosphingolipid biosynthesis - ganglio series | 0.0152 |
| path:mmu05202 Transcriptional misregulation in cancer | 0.0185 |
| path:mmu04066 HIF-1 signaling pathway | 0.0282 |
| path:mmu04080 Neuroactive ligand-receptor interaction | 0.0282 |
| path:mmu04350 TGF-beta signaling pathway | 0.0282 |
| path:mmu04917 Prolactin signaling pathway | 0.0422 |

## References

1. Bernstein, B.E. et al. A bivalent chromatin structure marks key developmental genes in embryonic stem cells. Cell 125, 315–26 (2006).

2. Jaenisch, R. & Young, R. Stem cells, the molecular circuitry of pluripotency and nuclear reprogramming. Cell 132, 567–82 (2008).

3. Zhou, Y., Kim, J., Yuan, X. & Braun, T. Epigenetic modifications of stem cells: a paradigm for the control of cardiac progenitor cells. Circ Res 109, 1067–81 (2011).

4. Boyer, L.A. et al. Polycomb complexes repress developmental regulators in murine embryonic stem cells. Nature 441, 349–53 (2006).

5. Brookes, E. et al. Polycomb associates genome-wide with a specific RNA polymerase II variant, and regulates metabolic genes in ESCs. Cell Stem Cell 10,157–70 (2012).

6. Hirose, Y. & Ohkuma, Y. Phosphorylation of the C-terminal domain of RNA polymerase II plays central roles in the integrated events of eucaryotic gene expression. J Biochem 141, 601–8 (2007).

7. Ferrai, C. et al. Poised transcription factories prime silent uPA gene prior to activation. PLoS Biol 8, e1000270 (2010).

8. Stock, J.K. et al. Ring1-mediated ubiquitination of H2A restrains poised RNA polymerase II at bivalent genes in mouse ES cells. Nat Cell Biol 9, 1428–35 (2007).

9. Akhtar, M.S. et al. TFIIH kinase places bivalent marks on the carboxy-terminal domain of RNA polymerase II. Mol Cell 34, 387–93 (2009).

10. Shema, E. et al. Single-molecule decoding of combinatorially modified nucleosomes. Science 352, 717–21 (2016).

11. Kolodziejczyk, A.A. et al. Single Cell RNA-Sequencing of Pluripotent States Unlocks Modular Transcriptional Variation. Cell Stem Cell 17, 471–85 (2015).

12. Tee, W.W., Shen, S.S., Oksuz, O., Narendra, V. & Reinberg, D. Erk1/2 activity promotes chromatin features and RNAPII phosphorylation at developmental promoters in mouse ESCs. Cell 156, 678–90 (2014).

13. Buettner, F. et al. Computational analysis of cell-to-cell heterogeneity in single-cell RNA-sequencing data reveals hidden subpopulations of cells. Nat Biotechnol 33, 155–60 (2015).

14. Islam, S. et al. Characterization of the single-cell transcriptional landscape by highly multiplex RNA-seq. Genome Res 21, 1160–7 (2011).

15. Tang, F. et al. mRNA-Seq whole-transcriptome analysis of a single cell. Nat Methods 6, 377–82 (2009).

16. Kim, J.K. & Marioni, J.C. Inferring the kinetics of stochastic gene expression from single-cell RNA-sequencing data. Genome Biol 14, R7 (2013).

17. Mikkelsen, T.S. et al. Genome-wide maps of chromatin state in pluripotent and lineage-committed cells. Nature 448, 553–60 (2007).

18. Brinkman, A.B. et al. Sequential ChIP-bisulfite sequencing enables direct genome-scale investigation of chromatin and DNA methylation cross-talk. Genome Res 22, 1128–38 (2012).

19. Fouse, S.D. et al Promoter CpG methylation contributes to ES cell gene regulation in parallel with Oct4/Nanog, PcG complex, and histone H3 K4/K27 trimethylation. Cell Stem Cell 2, 160–9 (2008).

20. Wang, J., Wen, S., Symmans, W.F., Pusztai, L. & Coombes, K.R. The bimodality index: a criterion for discovering and ranking bimodal signatures from cancer gene expression profiling data. Cancer Inform 7, 199–216(2009).

21. Sharova, L.V. et al. Database for mRNA half-life of 19 977 genes obtained by DNA microarray analysis of pluripotent and differentiating mouse embryonic stem cells. DNA Res 16, 45–58 (2009).

22. Niwa, H., Ogawa, K., Shimosato, D. & Adachi, K. A parallel circuit of LIF signalling pathways maintains pluripotency of mouse ES cells. Nature 460, 118–22 (2009).

23. Batada, N.N. & Hurst, L.D. Evolution of chromosome organization driven by selection for reduced gene expression noise. Nat Genet 39, 945–9 (2007).

24. Ebisuya, M., Yamamoto, T., Nakajima, M. & Nishida, E. Ripples from neighbouring transcription. Nat Cell Biol 10, 1106–13 (2008).

25. Hebenstreit, D., Deonarine, A., Babu, M.M. & Teichmann, S.A. Duel of the fates: the role of transcriptional circuits and noise in CD4+ cells. Curr Opin Cell Biol 24, 350–8 (2012).

26. Wang, G.Z., Lercher, M.J. & Hurst, L.D. Transcriptional coupling of neighboring genes and gene expression noise: evidence that gene orientation and noncoding transcripts are modulators of noise. Genome Biol Evol 3, 320–31 (2011).

27. Schoenfelder, S. et al. Polycomb repressive complex PRC1 spatially constrains the mouse embryonic stem cell genome. Nat Genet 47, 1179–86 (2015).

28. Schoenfelder, S. et al. The pluripotent regulatory circuitry connecting promoters to their long-range interacting elements. Genome Res 25, 582–97 (2015).

29. Endoh, M. et al. Polycomb group proteins Ring1A/B are functionally linked to the core transcriptional regulatory circuitry to maintain ES cell identity. Development 135, 1513–24 (2008).

30. Suganuma, T. & Workman, J.L. Signals and combinatorial functions of histone modifications. Annu Rev Biochem 80, 473–99 (2011).

31. Munsky, B., Neuert, G. & van Oudenaarden, A. Using gene expression noise to understand gene regulation. Science 336, 183–7 (2012).

32. Raj, A. & van Oudenaarden, A. Single-molecule approaches to stochastic gene expression. Annu Rev Biophys 38, 255–70 (2009).

33. Barkai, N. & Shilo, B.Z. Variability and robustness in biomolecular systems. Mol Cell 28, 755–60 (2007).

34. Bar-Even, A. et al. Noise in protein expression scales with natural protein abundance. Nat Genet 38, 636–43 (2006).

35. Elowitz, M.B., Levine, A.J., Siggia, E.D. & Swain, P.S. Stochastic gene expression in a single cell. Science 297, 1183–6 (2002).

36. Weinberger, L. et al. Expression noise and acetylation profiles distinguish HDAC functions. Mol Cell 47, 193–202 (2012).

37. Eldar, A. & Elowitz, M.B. Functional roles for noise in genetic circuits. Nature 467, 167–73 (2010).

38. Kumar, R.M. et al. Deconstructing transcriptional heterogeneity in pluripotent stem cells. Nature 516, 56–61 (2014).

39. Cloonan, N. et al. Stem cell transcriptome profiling via massive-scale mRNA sequencing. Nat Methods 5, 613–9 (2008).

40. Nishiyama, A. et al. Uncovering early response of gene regulatory networks in ESCs by systematic induction of transcription factors. Cell Stem Cell 5, 420–33 (2009).

41. Young, M.D. et al. ChIP-seq analysis reveals distinct H3K27me3 profiles that correlate with transcriptional activity. Nucleic Acids Res 39, 7415–27 (2011).

42. Sanchez, A., Choubey, S. & Kondev, J. Regulation of noise in gene expression. Annu Rev Biophys 42, 469–91 (2013).

43. Batenchuk, C. et al. Chromosomal position effects are linked to sir2-mediated variation in transcriptional burst size. Biophys J 100, L56–8 (2011).

44. Becskei, A., Kaufmann, B.B. & van Oudenaarden, A. Contributions of low molecule number and chromosomal positioning to stochastic gene expression. Nat Genet 37, 937–44 (2005).

45. Dar, R.D. et al. Transcriptional burst frequency and burst size are equally modulated across the human genome. Proc Natl Acad Sci U S A 109, 17454–9 (2012).

46. Raj, A., Peskin, C.S., Tranchina, D., Vargas, D.Y. & Tyagi, S. Stochastic mRNA synthesis in mammalian cells. PLoS Biol 4, e309 (2006).

47. Skupsky, R., Burnett, J.C., Foley, J.E., Schaffer, D.V. & Arkin, A.P. HIV promoter integration site primarily modulates transcriptional burst size rather than frequency. PLoS Comput Biol 6 (2010).

48. Suter, D.M. et al. Mammalian genes are transcribed with widely different bursting kinetics. Science 332, 472–4 (2011).

49. Newman, J.R. et al. Single-cell proteomic analysis of S. cerevisiae reveals the architecture of biological noise. Nature 441, 840–6 (2006).

50. Bulger, M. & Groudine, M. Functional and mechanistic diversity of distal transcription enhancers. Cell 144, 327–39 (2011).

51. Wu, T.D. & Nacu, S. Fast and SNP-tolerant detection of complex variants and splicing in short reads. Bioinformatics 26, 873–81 (2010).

52. Anders, S. & Huber, W. Differential expression analysis for sequence count data. Genome Biol 11, R106 (2010).

53. Smedley, D. et al. The BioMart community portal: an innovative alternative to large, centralized data repositories. Nucleic Acids Res 43, W589–98 (2015).

54. Langmead, B. & Salzberg, S.L. Fast gapped-read alignment with Bowtie 2. Nat Methods 9, 357–9 (2012).

55. Xing, H., Mo, Y., Liao, W. & Zhang, M.Q. Genome-wide localization of protein-DNA binding and histone modification by a Bayesian change-point method with ChIP-seq data. PLoS Comput Biol 8, e1002613 (2012).

56. Singer, G.A., Lloyd, A.T., Huminiecki, L.B. & Wolfe, K.H. Clusters of coexpressed genes in mammalian genomes are conserved by natural selection. Mol Biol Evol 22, 767–75 (2005).

57. Kanehisa, M. & Goto, S. KEGG: kyoto encyclopedia of genes and genomes. Nucleic Acids Res 28, 27–30 (2000).

